# Ccube: A fast and robust method for estimating cancer cell fractions

**DOI:** 10.1101/484402

**Authors:** Ke Yuan, Geoff Macintyre, Wei Liu, PCAWG-11 working group, Florian Markowetz

**Author notes:** Correspondence to: Ke Yuan and Florian Markowetz.

## Abstract

Estimating and clustering cancer cell fractions of genomic alterations are central tasks for studying intratumour heterogeneity. We present Ccube, a probabilistic framework for inferring the cancer cell fraction of somatic point mutations and the subclonal composition from whole-genome sequencing data. We develop a variational inference method for model fitting, which allows us to handle samples with large number of the variants (more than 2 million) while quantifying uncertainty in a Bayesian fashion. Ccube is available at https://github.com/keyuan/ccube.

## 1. Introduction

A fundamental problem when studying intratumour heterogeneity is to estimate the cancer cell fraction (CCF) of a single nucleotide variants (SNVs). The key difficulty is that CCF is proportional to the number of mutated chromosomal copies, known as the multiplicity of a mutation, which is also unknown. For any given mutation, the observed variant allele frequency can be modelled by CCF times multiplicity. As a result, it is impossible to estimate CCF without making strong assumptions about what multiplicity is. Several methods choose to prefix multiplicities, for example, DPClust [1] and PyClone [2]. PhyloSub [3] and PhyloWGS [4] use phylogenetic trees to estimate the multiplicities. This approach significantly increase the complexity of the model making model inference difficult to scale.

We develop Ccube, a method using clustering (i.e. assuming multiple mutations share the same CCF) to determine what the appropriate multiplicities are. The method takes sequencing reads profiles of SNVs, corrects them for copy number alterations and purity, and produces CCF estimates for all mutations within the sample.

## 2. Results

### 2.1. Mapping between variant allele frequency and cancer cell fraction

Given the purity of the sample and copy number profile at the locus of a SNV of interest, there is a mapping between VAF and the CCF. Following [2], we formulate relationship between VAF and CCF from a probability stand point. Generally speaking, we are dealing with two questions here: first, are we observing a variant allele on the a sequencing read covering the locus of the mutation i.e. read variable. Second, where is the read comes from i.e. population variable. The key for the probabilistic view point is to consider VAF as the marginal probability of observing a variant read, where the population variable is integrated out:

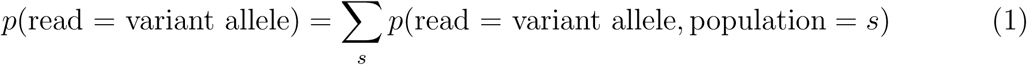

Following the definition in [2], the population variable has three possible states: normal cell population, cancer cells that don’t bear the SNV, defined as the reference population and cancer cells that carry the mutation, defined as the variant population. Based these three possible populations. For each population, we have the probability of observing a read coming from the population, and the conditional probability of the population contributing a variant read. The marginalisation can be written as,

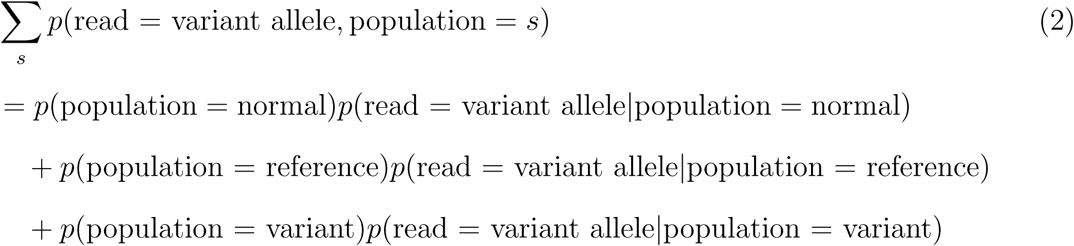

The probability of observing a read coming from a population is proportional to the prevalence of the population in the sample times its total copy number. Specifically, the prevalence the variant population is CCF, *ϕ*. Therefore, we have

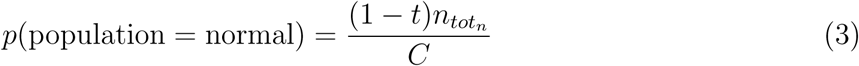

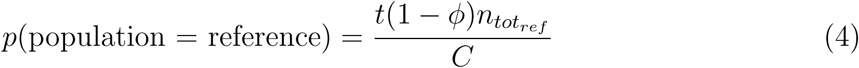

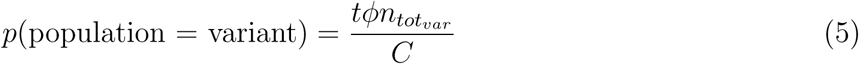

where 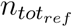, 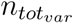, 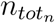 are the total copy numbers of cells in the reference, variant, and normal populations, respectively. The normalising constant, 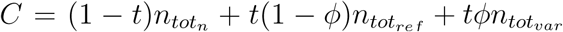, make sure the probabilities add up to one.

We further assume the reference shares the same total copy number with the variant population,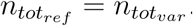. This corresponds to assuming all the cancer cells in the sample share the same total copy number at the site of SNV, i.e. the copy number is clonal [1]. Using 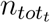 to represent the clonal total copy number. We have:

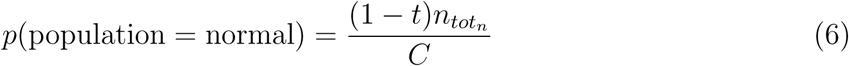

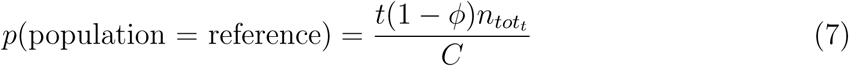

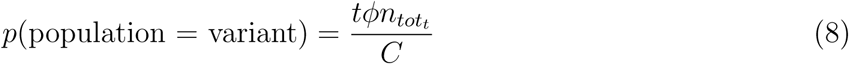

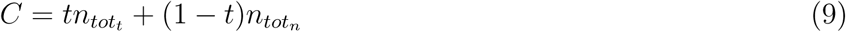

The conditional probabilities for populations to produce a variant read are:

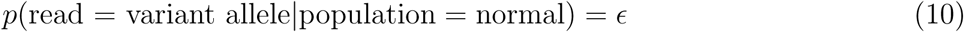

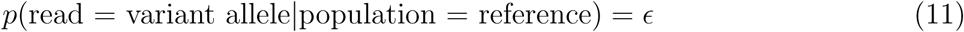

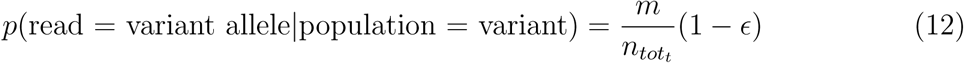

where *ϵ* is a uniform sequencing error. *m* is the number of mutated chromosomal copy, the multiplicity of the mutations.

Taken together, we obtain a linear mapping between the probability of observing a variant read at a mutated locus, *f,* and the CCF of the mutation *ϕ*:

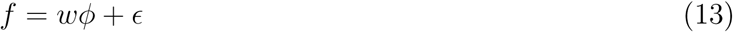

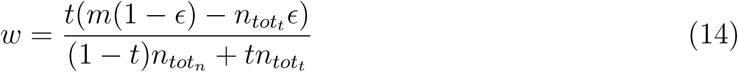

Assuming the number of read carrying the variant allele follows a binomial distribution, the VAF is an unbiased estimator of *f*.

### 2.2. The Ccube model for estimating and clustering cancer cell fractions

Let *i ∈ {*1*,…, N}* denotes the index for each variant considered, and *k ∈* 1*,…, K* denote the index for the number of CCF cluster identified in the mixture. For the *i*th variant, *b*_*i*_ and *d*_*i*_ denote the number of reads reporting variant allele and the total number of reads. The copy number profile at the *i*th mutation includes the total copy number of the tumour 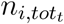, the total copy number of the normal population 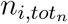, the copy number of the major allele 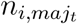, the copy number of the minor allele 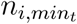 The number of mutated chromosomal copies is denoted by *m*_*i*_.

The basis of Ccube is a Binomial mixture model. We assume the *i*th variant allele read count *b*_*i*_ follows a Binomial Distribution with total read count *d*_*i*_ and expected VAF *f*_*i*_ as its parameter

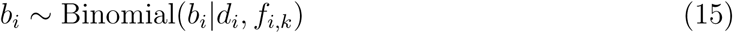

Where

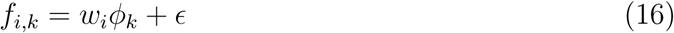

. where 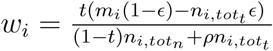.

The Ccube model can be shown as the following:

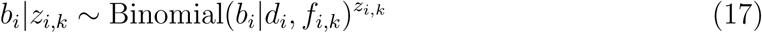

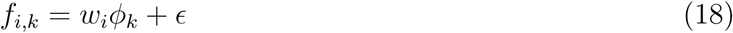

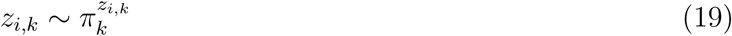

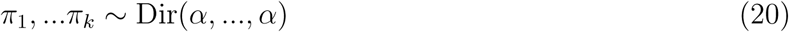

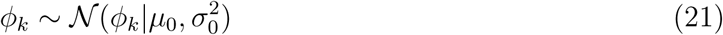

### 2.3. Variational inference for Ccube

The variational inference maximises the evidence lower bound (ELBO) of the marginal likelihood of the model:

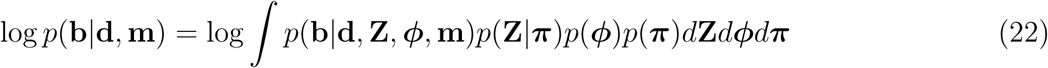

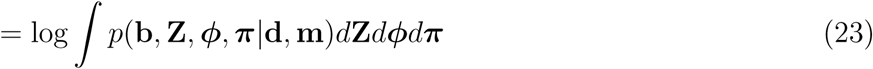

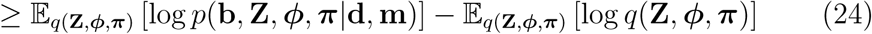

where **b** = *{b*_*i*_*}*, **d** = *{d*_*i*_*}*, **Z** = *{z*_*i,k*_*}*, ***ϕ*** = *{ϕ*_*k*_*}*, ***π*** = *{π*_*k*_*}*, **m** = *{m*_*i*_*}*.

We adopt the common fixed-form mean field approximation, in which **Z**, ***ϕ***, ***π*** are independent:

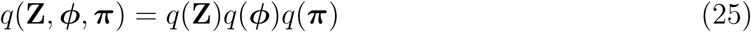

Maximising the ELBO with respect to the above *q*(**Z**, ***ϕ***, ***π***) yields the following forms:

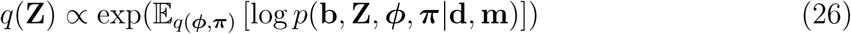

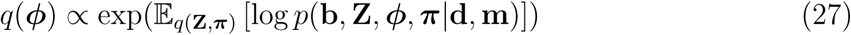

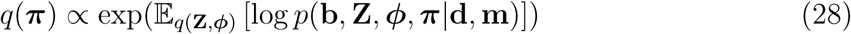

These distributions are fitted to the data with a variational Ε-step and variational M-step. In the Ε-step, we update the approximate posterior of the assignment.

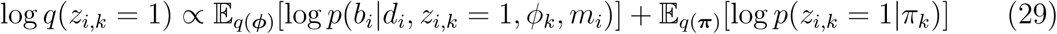

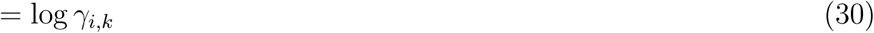

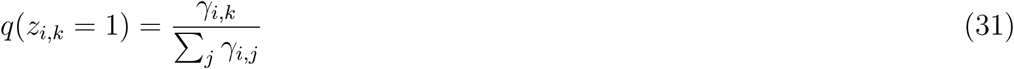

In the variational M-step, we update the approximate posteriors on parameters ***ϕ*** and ***π***.

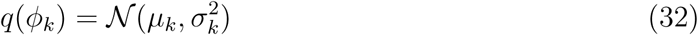

where,

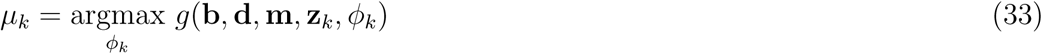

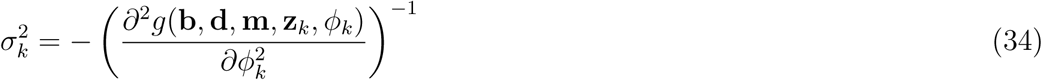

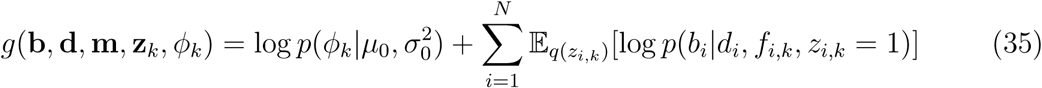

The multiplicities are estimated as the following

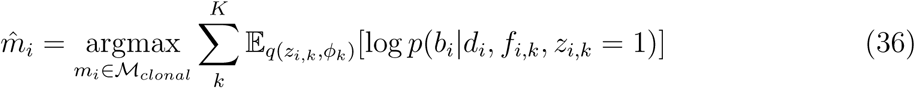

where 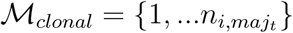.

Finally, *q*(***π***) is obtained as standard variational approximation for mixing weight in mixture models [5].

## 3. Ccube pipeline

Preprocessing: The Ccube pipeline uses the clonal copy number, consensus purity and variant and reference allele read counts from somatic point mutation calls.

Postprocessing steps: The core variational inference is ran with a range of possible number of clusters. The solution with the best ELBO is selected. The solution is consisted of 1) posterior distributions of in terms of means and variances; 2) posterior probabilities of each mutation to be assigned to all clusters and final assignments; 3) multiplicities and observed CCFs based multiplicities. Clusters with less than 1% of mutation assigned are removed. The mutations are re-assigned with variational expectation step. Clusters with mean CCF closer than 10% are merged by re-running the inference with merged cluster configuration. A typical graphical summary can be found in figure 1.

**Figure 1:**
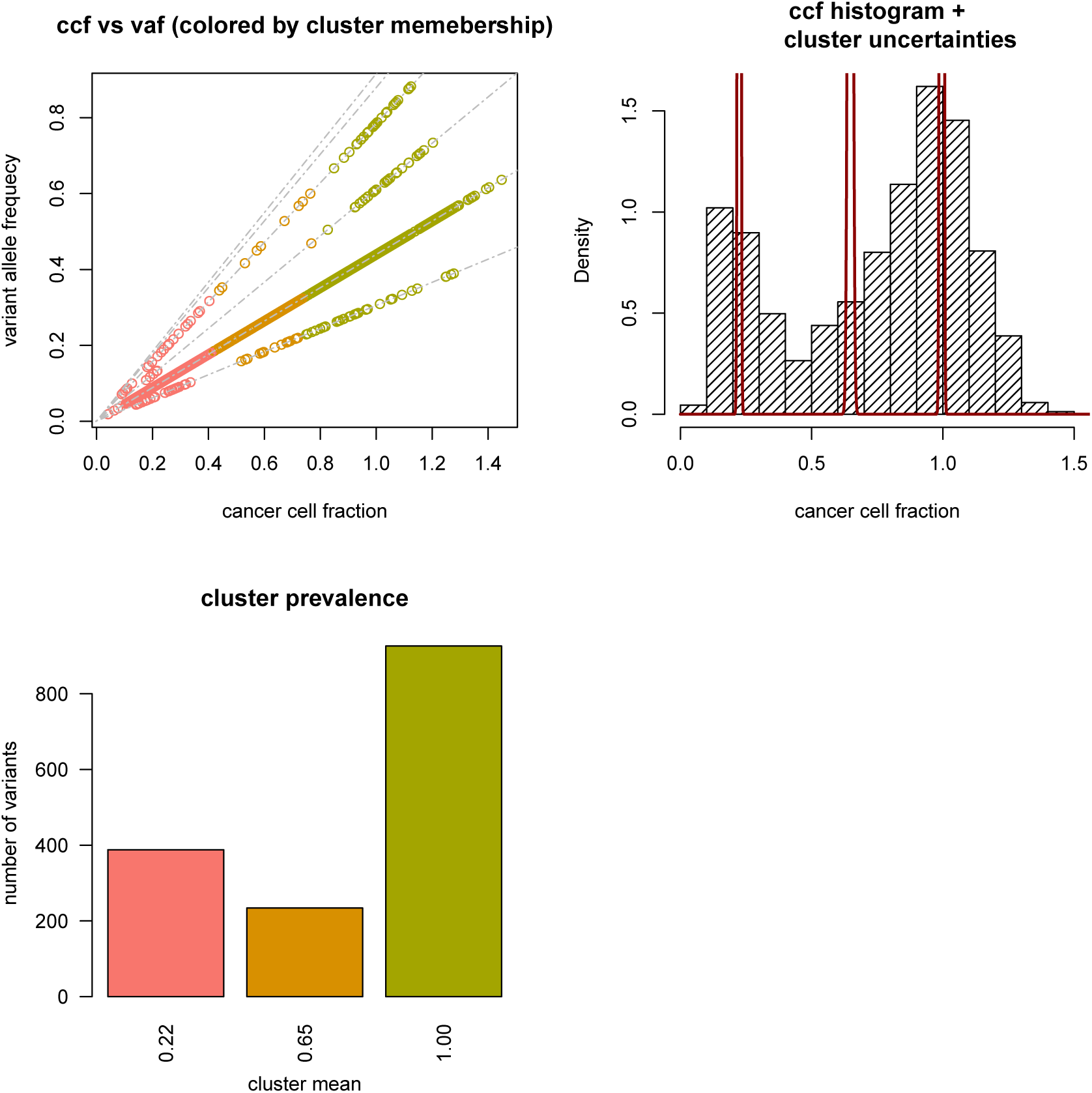
An example of Ccube sample results summary. A: Scatter plot of VAF and CCF. Each point in the figure is a mutation color coded by its cluster membership. The gray dashed line are all possible linear mappings (eq. 1) determined by copy number and multiplicity configurations in the sample. B: Histogram of observed CCFs. The red solid line shows the approximated posterior distribution of CCF cluster centers. The peak at CCF=1 corresponds to the clonal cluster. C: The number of variants assigned each CCF cluster. Each CCF cluster is labelled by it cluster center.

### 3.1. Estimating purity

Ccube pipeline also produces an independent purity estimate using mutations from balanced copy number regions. For samples without whole-genome duplication, only mutations in normal copy number regions are included. For samples with whole-genome duplication, all mutations in balanced copy number regions are included. We then convert the allele frequencies to cellular prevalence using the following equation:

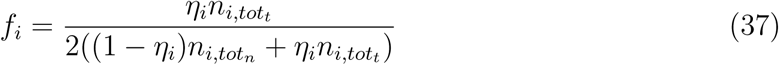

Where is the VAF of the *i*th mutation obtained as the ratio between variant and wild-type allele counts, *η* is the cellular prevalence of the th mutation, are the total copy number of normal and tumour populations respectively. We then cluster the using students-t mixture model. The model is fitted with the variational Bayes approach described in [6]. The purity corresponds to the component with the largest mean, in additional the eligible component must have at least more than 1.5% of mutation assigned to it.

